# DNA-PAINT imaging with hydrogel imprinting and clearing

**DOI:** 10.1101/2025.02.14.638341

**Authors:** Johannes Stein, Lorenzo Magni, George M. Church

**Affiliations:** Wyss Institute of Biologically Inspired Engineering, Boston, MA, USA; Department of Genetics, Harvard Medical School, Boston, MA, USA

## Abstract

Hydrogel-embedding is a versatile technique in fluorescence microscopy, offering stabilization, optical clearing, and the physical expansion of biological specimens. DNA-PAINT is a super-resolution microscopy approach based on the diffusion and transient binding of fluorescently labeled oligos, but its feasibility in hydrogels has not yet been explored. In this study, we demonstrate that polyacrylamide hydrogels support sufficient diffusion for effective DNA-PAINT imaging. Using acrydite-anchored oligonucleotides imprinted from patterned DNA origami nanostructures and microtubule filaments in fixed cells, we find that hydrogel embedding preserves docking strand positioning at the nanoscale. Sample clearing via protease treatment had minor structural effects on the microtubule structure, and enhanced diffusion and accessibility to hydrogel-imprinted docking strands. Our work demonstrates promising potential for diffusion and binding-based fluorescence imaging applications in hydrogel-embedded samples.

**Graphical abstract:** 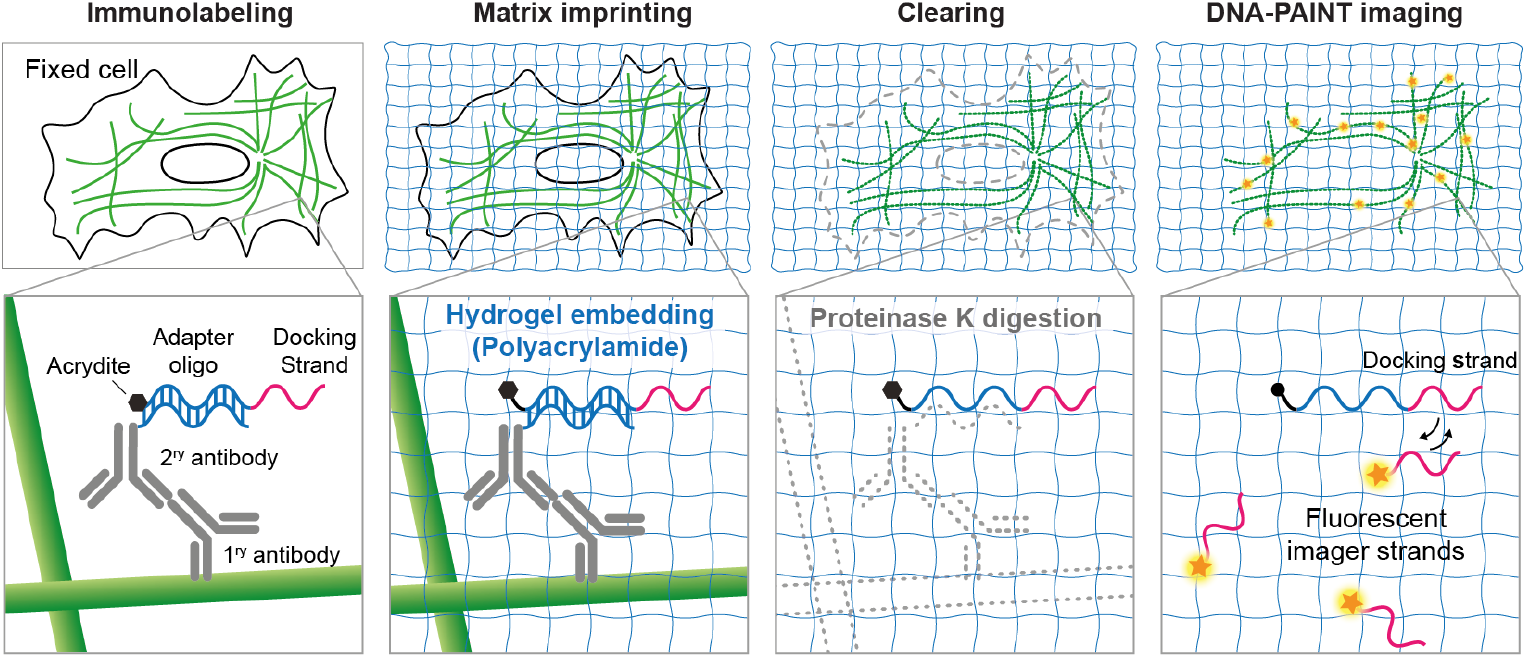

## Introduction

Hydrogels can enhance fluorescence microscopy applications across diverse biological systems. Their ability to stabilize, clear, and render samples optically transparent has significantly advanced imaging workflows of larger samples such as tissues, organs and even entire mammalian bodies^1–3^. Moreover, hydrogels enable super-resolution (SR) imaging on conventional light microscopes by leveraging swellable polymers to physically expand biological samples – a technique known as Expansion Microscopy^4,5^ (ExM).

Optical SR approaches such as STED^6^ (Stimulated Emission Depletion), SIM^7,8^ (Structured Illumination Microscopy), PALM^9^ (Photo-Activated Localization Microscopy) or STORM^10^ (Stochastic Optical Reconstruction Microscopy) offer significant potential when combined with hydrogel embedding. For instance, hybrid implementations can benefit from sample clearing, enhanced resolution in ExM, and validation of pre- and post-expansion morphologies^11–18^. To prevent loss of spatial information during sample clearing or proteolytic homogenization required for ExM, DNA-based secondary probes have been employed to imprint the positions of target proteins or nucleic acids into the hydrogel matrix^4,19–24^. This DNA-anchoring strategy is particularly advantageous for multiplexing approaches such as sequential barcoding and in situ sequencing^20,23–25^. In this context, the removal of endogenous cellular components has been shown to enhance target specificity and signal-to-noise ratios in “matrix-imprinted” hydrogels^20,23^. Notably, prolonged hybridization and washing steps have proven sufficient for fluorescently labeled reporter oligonucleotides to overcome the reduced diffusion properties inherent to hydrogels^26^.

DNA-PAINT^27^ (Points Accumulation for Imaging in Nanoscale Topography) is a SR imaging technique that relies on the transient binding of short, dye-labeled “imager” oligonucleotides to complementary “docking strands” attached to target molecules. This dynamic binding mechanism generates stochastic single-molecule fluorescence ‘blinking’, enabling SR imaging with sub-10 nm resolution^28^, resistance to photobleaching^29^, and spectrally-unlimited multiplexing^30^. However, DNA-PAINT depends on sufficient diffusion and binding turnover of imager strands inside the sample. While single dye molecules have been shown to diffuse effectively for direct PAINT imaging of synthetic hydrogel structures^31^, the compatibility of DNA-PAINT – typically using 7–10 nucleotide-long imager strands – with hydrogel embedding remains unexplored.

Here, we demonstrate that DNA-PAINT imaging can be effectively applied to biological samples embedded in polyacrylamide (PA) hydrogels. Building on previous adaptations of target-imprinted DNA origami nanostructures for STORM imaging^32^, we validate that hydrogel embedding does not significantly affect sample structure or achievable resolution. Furthermore, we find that reduced imager binding rates due to restricted diffusion can be compensated by increasing imager strand concentrations. While proteolytic degradation for clearing introduced nanoscale distortions of in vitro DNA origami patterns, microtubule networks in fixed cells remained structurally intact, likely due to the stabilizing properties of the 3D cellular hydrogel matrix. Our findings highlight promising potential for hydrogel-based DNA-PAINT applications and serve as a useful starting point for future applications in expanded hydrogels.

## Results

### Feasibility and in vitro assessment of DNA-PAINT with hydrogel matrix imprinting and clearing

To assess the compatibility of DNA-PAINT with hydrogel embedding, we adapted an approach to incorporate oligonucleotide adapters hybridized to patterned DNA origami nanostructures into hydrogels^32^, allowing to probe possible structural deformations introduced by the embedding process. We selected a well-established DNA origami design featuring 3×4 adapter binding sites arranged at a distance of 20 nm^29^. DNA origami structures were immobilized on the surface of open imaging chambers and adapter strands were added. These strands featured a single-stranded overhang serving as a docking site for DNA-PAINT on the 5′ end and an acrydite moiety on the 3′ end, ensuring that the latter was positioned close to the origami surface upon hybridization (**Fig. 1**a). To validate the functionality of this design prior to hydrogel embedding we performed standard DNA-PAINT imaging using Total Internal Reflection Fluorescence (TIRF) microscopy. We achieved a localization precision of ∼4.2 nm (determined via Nearest Neighbor Analysis^33^; σ_NeNA_) and the fully resolved 20 nm pattern in averaged origami sum images confirmed that the acrydite modification did not negatively impact imaging performance (**Fig. 1**b; see **Supplementary Fig. 1** for a random selection of 400 individual DNA origami structures).

Next, we polymerized polyacrylamide (PA) hydrogels within the same imaging chambers to evaluate the feasibility of DNA-PAINT imaging inside hydrogels and assess potential effects on structural integrity and achievable resolution. Our initial observation was that, whereas imagers typically arrive instantaneously under standard aqueous conditions after the addition of mixed imaging buffers, the slower diffusion within hydrogels caused a ∼30 s delay before the first single-molecule blinking events were observed at the glass surface. Even after equilibration for 30 min, the blinking density remained significantly lower compared to standard DNA-PAINT at the same imager concentration. To compensate for this reduced imager association rate, we increased imager concentrations by approximately 4-fold, which restored optimal blinking densities. TIRF imaging under these conditions successfully resolved imprinted DNA origami patterns, confirming the feasibility of DNA-PAINT imaging within PA hydrogels (**Fig. 1**c and **Supplementary Fig. 2**). Since we observed comparable numbers of docking strands per origami before and after embedding, the reduced blinking density suggests a ∼4× decrease in imager association rate, which can, however, be counteracted by increasing imager concentrations. While this adjustment may elevate background fluorescence levels and affect signal-to-noise ratios, we observed only a mild reduction in localization precision (∼5.7 nm).

Since hydrogel clearing protocols for biological samples often involve proteolytic digestion, we sought to evaluate the impact of such treatments using our DNA origami samples. We subjected hydrogel-embedded DNA origami structures to Proteinase K digestion (1 h at 37°C), a standard procedure used to degrade cellular proteins and reduce non-specific interactions of fluorescent readout oligonucleotides in multiplexed transcriptomics applications^23^. However, we observed a notable reduction in resolution following digestion (**Fig. 1**d and **Supplementary Fig. 3**). Averaging was only feasible for manually selected origami structures, suggesting that clearing-induced nanoscale distortions disrupted the imprinted pattern (**Supplementary Fig. 4**). Since Proteinase K should not degrade DNA or the polyacrylamide gel itself, it is likely that the elevated temperature and low ion content of the digestion buffer contributed to destabilization of both the DNA origami and the hydrogel matrix. To quantitatively assess whether the positioning of docking strands followed the expected spatial distribution, we leveraged the known rectangular 20 nm grid pattern of our DNA origami structures. Using DBSCAN clustering, we identified individual docking strands and computed the center-of-mass distances to the five nearest docking strands (**Supplementary Fig. 5**). **Fig. 1**e displays the nearest-neighbor distance distributions for standard DNA-PAINT imaging, hydrogel-embedded samples, and hydrogel-embedded samples subjected to clearing. The blue dashed lines indicate the expected peak positions for an ideal 20 nm rectangular grid (see also **Supplementary Fig. 5**). Despite the reduced resolution affecting peak width in hydrogel-embedded samples, the overall nearestneighbor distribution remained similar to that of unembedded origami, as had been previously observed for PA hydrogel imprinting using STORM^32^. However, incubation in clearing buffer resulted in nanoscale distortions, leading to an average 25% increase in the nearest-neighbor distance, with the first peak shifting to ∼25 nm.

### DNA-PAINT imaging of hydrogel-imprinted and cleared cells

Encouraged by our *in vitro* results, we next investigated the feasibility of hydrogel embedding and clearing for DNA-PAINT imaging in biological specimens. We labeled α-tubulin in fixed HEK293T cells using primary antibodies and secondary antibodies carrying the same adapter binding site previously used for DNA origami nanostructures. As a first step, we performed standard DNA-PAINT imaging without hydrogel embedding to confirm robust hybridization of docking strand adapters to secondary antibodies and ensure compatibility with cellular imaging (**Fig. 2b**).

**Figure 2.**
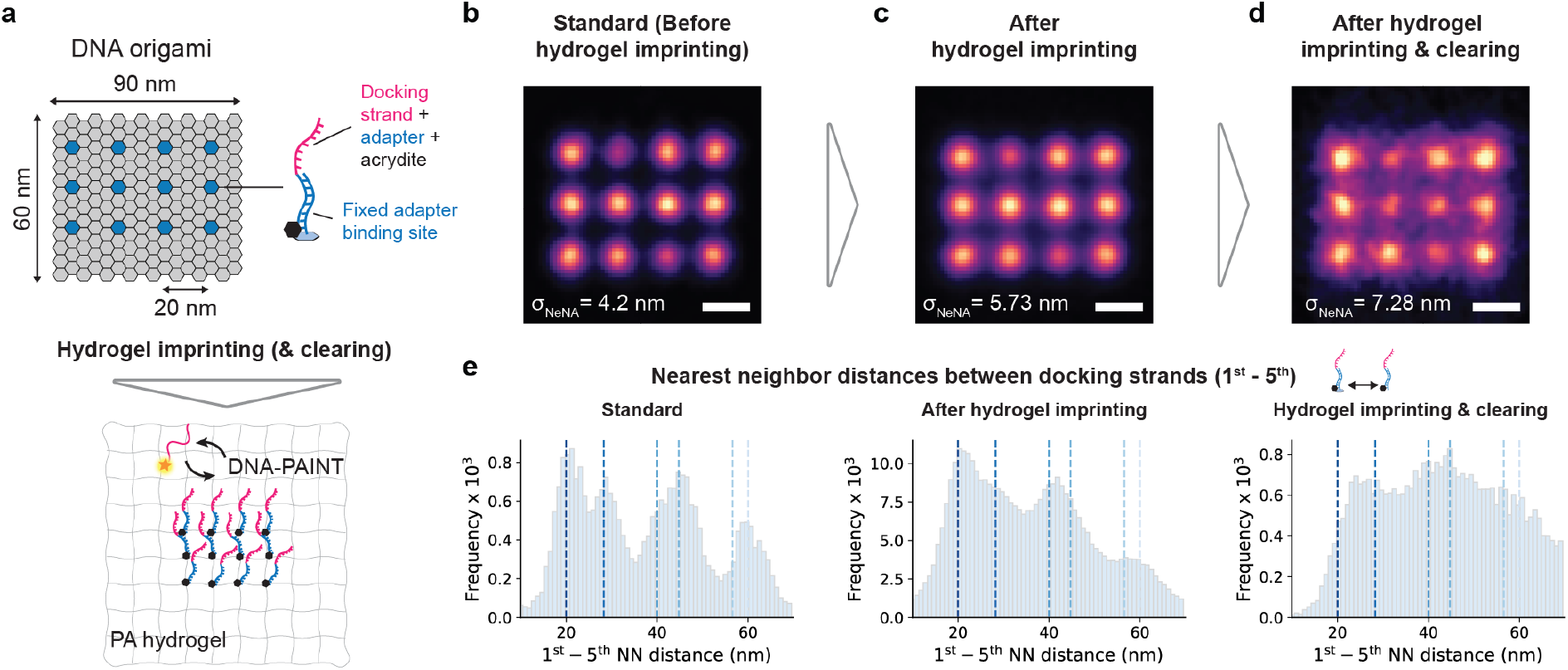
Feasibility and in vitro assessment of DNA-PAINT with hydrogel matrix imprinting. (a) Experimental scheme inspired by Lee *et al*.^32^ for testing DNA-PAINT imaging in hydrogels using synthetic DNA origami structures. Origami surfaces featured a 3×4 pattern of binding sites for acrydite-labeled docking strand adapter oligonucleotides arranged at a 20 nm spacing. Upon polymerization of PA, acrydite is incorporated into the hydrogel, imprinting the docking strand pattern for subsequent DNA-PAINT imaging (n=1,951). (b) Averaged DNA-PAINT sum image 20 nm docking strand pattern without hydrogel embedding (n=379). (c) Averaged DNA-PAINT sum image 20 nm docking strand pattern after hydrogel embedding (n=379). (d) Averaged DNA-PAINT sum image 20 nm docking strand pattern after hydrogel embedding and incubation with Proteinase K, commonly used to degrade cellular protein. (n=159; hand-selected). (e) Profiles of nearest-neighbor distances between docking strands for DNA origami datasets in (b-d). The distributions contain the distances of each docking strand to its 5 nearest neighbors and the expected distances from the 20 nm grid pattern are indicated by blue dashed lines (see also Supplementary Fig. 4). Scale bars, 20 nm.

Using the same PA hydrogel conditions previously used in vitro, we noticed that during washing steps hydrogels could detach from the surfaces and observed increased non-specific imager sticking at the glass-hydrogel. To mitigate these issues, we coated cover glasses with polylysine, which improved hydrogel adherence to the glass and cell spreading for TIRF imaging. This was further aided by polymerizing thinner PA gels as previously explored for ExM^34,35^. Thinner hydrogels had the additional advantage of reducing the lag time between addition of the imaging buffer and equilibration of the blinking density.

Given that hydrogel embedding without proteolytic degradation increases intracellular density, we anticipated that DNA-PAINT imaging might be hindered due to limited imager strand access. Surprisingly, the presence of the PA hydrogel did not significantly alter diffusion conditions. As observed with DNA origami samples, a ∼4-fold increase in imager concentration – compared to standard DNA-PAINT of unembedded cells – allowed for effective imaging, with ∼20 min required for equilibration before acquisition. Under these conditions, we successfully reconstructed super-resolved images of the microtubule network in hydrogel-embedded cells (**Fig. 2c**).

To further explore the potential of hydrogel matrix imprinting and clearing, we treated hydrogel-embedded HEK cells with Proteinase K to degrade endogenous proteins, generating both imprinted and cleared hydrogels. Since PA does not interact with endogenous cellular components but forms a surrounding hydrogel matrix, we hypothesized that proteolysis would enhance imager diffusion and improve docking strand accessibility. Indeed, we found that imager concentrations could be reduced by twofold while still maintaining optimal single-molecule blinking. However, diffusion remained significantly slower than in unembedded cells, still necessitating short equilibration periods (∼10 min after imager strand addition) before imaging. **Fig. 2d** shows an exemplary superresolved image of the imprinted microtubule network of a cleared cell, demonstrating the feasibility of integrating hydrogel matrix imprinting and clearing with DNA-PAINT.

To assess imaging performance with respect to resolution, we computed the localization precision (σ_NeNA_) for three data sets acquired at each conditions: standard DNA-PAINT (purple), hydrogel-imprinted (red), and hydrogel-embedded & cleared (yellow) (**Fig. 2e**). While localization precisions varied within experimental limits, we observed a slight reduction in precision when comparing standard DNA-PAINT to hydrogel-imprinted samples (∼6.1 nm vs. ∼7.4 nm, respectively), likely due to increased background fluorescence from higher imager concentrations. Interestingly, the reduction in imager concentration enabled by clearing improved localization precision (∼6.5 nm), approaching that of unembedded samples. Since σ_NeNA_ estimates the localization uncertainty of single emitters (in case of DNA-PAINT a single docking strand), this implies that after proteolysis imprinted docking strands either maintain their positions to a similar extent as in unembedded cells or possible motility of imprinted docking strands is beyond our achievable resolution.

Finally, we assessed potential structural effects by measuring the average diameter of individual microtubules across experimental conditions (**Fig. 2f**; 15 microtubule cross-sections per condition). Consistent with our DNA origami findings, hydrogel embedding alone resulted in microtubule widths comparable to those of unembedded samples (∼50 nm). Note that indirect immunolabeling introduces a ∼25 nm linkage error due to antibody size, meaning that these measurements align with expected values. As previously demonstrated for ExM^4^, microtubule diameters are relatively robust to hydrogel embedding, enzymatic digestion, and expansion. In our case, Proteinase K treatment led to a moderate broadening of the microtubule diameter (∼54 nm), though the difference was not statistically significant. However, the increased variability in diameter measurements suggests minor structural distortions, which could potentially be mitigated through hydrogel protocols specifically optimized for ultrastructural preservation^36,37^.

**Figure 2.**
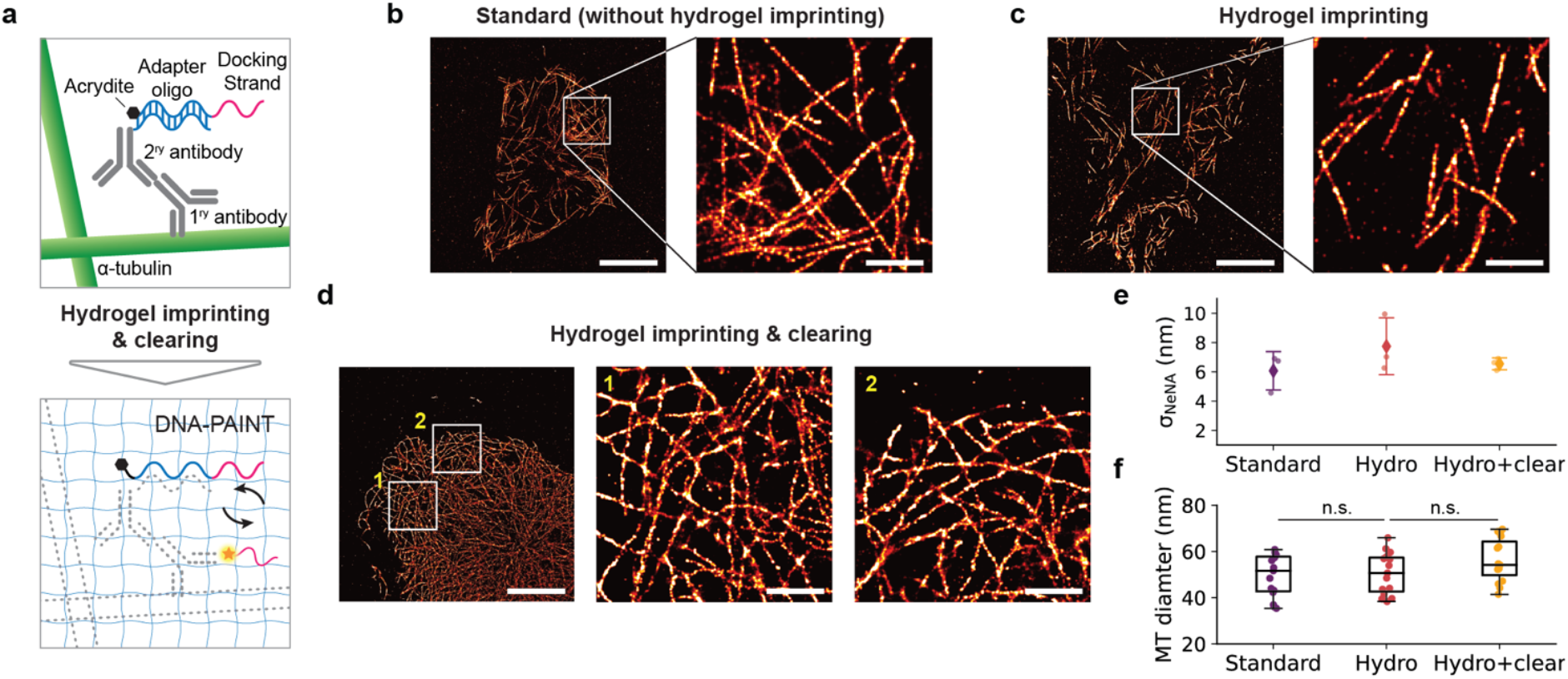
DNA-PAINT imaging of hydrogel-imprinted and cleared cells. (a) Schematic of the hydrogel imprinting and clearing approach^23^. Fixed cells were labeled for α-tubulin using primary and secondary antibodies. Secondary antibodies carried binding sites for acrydite-labeled docking strand adapter oligonucleotides for imprinting into PA hydrogels. After embedding, enzymatic protein degradation is used to clear the imprinted hydrogel. (b) Reconstructed DNA-PAINT image of microtubule filaments acquired without hydrogel imprinting. (c) Reconstructed DNA-PAINT image of microtubule filaments acquired with hydrogel imprinting. (d) Reconstructed DNA-PAINT image of microtubule filaments acquired with hydrogel imprinting and clearing. (e) Localization precision σ_NeNA_ obtained for three data sets per experimental condition in (a-c). (f) Measured microtubule diameters across filaments (n=15 per condition). Scale bars, 10 µm and 2 µm in zoom-ins.

## Discussion

In this work, we demonstrated the compatibility of DNA-PAINT imaging with hydrogel matrix imprinting and clearing. Following common embedding schemes^4,32^, we leveraged oligonucleotide adapters containing a DNA-PAINT docking strand and an acrydite moiety for stable incorporation into PA hydrogels. Targeting DNA origami as patterned control structures^32^ and microtubule filaments in fixed cells, we found that hydrogel embedding can preserve docking strand positioning both *in vitro* and *in situ*. Increasing imager concentrations to balance reduced imager diffusion facilitated efficient DNA-PAINT imaging with only minor reduction in achievable resolution. Applying proteolytic clearing via treatment with Proteinase K led to distortion of the origami pattern, likely due to temperature and buffer induced loss of DNA origami structure at the hydrogel surface. As expected from earlier ExM applications, proteinase treatment of hydrogel-imprinted cells revealed overall structural preservation of microtubule filaments with minor broadening of immunolabeled microtubule diameters. The clearing process itself enhanced imager diffusion by providing a homogenized and more accessible matrix.

Future applications of hydrogel-based DNA-PAINT could benefit from steady flow-based fluidics, which has been shown to significantly enhance imager diffusion in thicker samples, such as tissues^38^. Additionally, background fluorescence could be minimized by implementing fluorogenic imagers that remain quenched during diffusion^39^. The homogenized matrix of cleared samples may also help reduce non-specific imager binding^33^, benefiting quantitative imaging approaches such as qPAINT^40^ (quantitative DNA-PAINT), localization-based Fluorescence Correlation Spectroscopy^41,42^ and kinetic multiplexing^43^.

Moreover, DNA-PAINT could be combined with ExM in an iterative embedding approach that prevents sample shrinkage caused by ionic buffers required for efficient DNA hybridization^25^. Such joint implementations could enable precise ultrastructural validations of the same target structures before and after expansion^36,37^. Furthermore, the large linkage error inherent to antibody-based labeling could potentially be reduced by pursuing post-expansion labeling strategies^44^. Finally, the controllable blinking dynamics of DNA-PAINT could benefit One-step Nanoscale ExM”^45^, a recent approach leveraging blinking-induced fluctuation analyses for structural studies, as well as strategies for protein fingerprinting^46,47^. In conclusion, our work provides a valuable foundation for future applications of DNA-PAINT in hydrogel-embedded samples.

## Author contribuGons

J.S. and L.M. contributed equally. J.S. performed experiments, analyzed data and wrote the manuscript. L.M. performed experiments, analyzed data and wrote the manuscript. J.S. and G.M.C. conceived the study. G.M.C. supervised the study. All authors read and approved the manuscript.

## Funding

J.S. acknowledges support by the European Molecular Biology Organization (ALTF 816-2021). L.M. acknowledges support by the Giovanni Armenise Harvard Foundation, the Ermenegildo Zegna Foundation, Scuola Normale Superiore and the Erasmus Program of University of Pisa. G.M.C. acknowledges support by the Department of Energy (DE-FG02-02ER63445). This work was also supported by NIH awards to Chao-ting Wu (5RM1HG011016-03). The MicRoN Imaging Facility at Harvard Medical School acknowledges support by NIH Grant S10 RR027344-01.

### Acknowledgements

We thank Takeyuki Miyawaki for helpful discussions and valuable experimental support regarding hydrogel embedding. We thank Jenny Tam for providing infrastructure and experimental support. We thank Benjamin Angulo for sharing reagents and Matthew Serrata, Michel Nofal and Ryan McMillan for support with DNA-antibody conjugation and helpful discussions. We acknowledge the MicRoN Imaging Facility at Harvard Medical School, where all imaging was performed, and thank Paula De La Milagrosa Montero Llopis, Praju Vikas Anekal and Adrienne Wells for experimental support. We thank Talley Lambert at the Nikon Imaging Center for helpful discussions and support. We thank Maurice Perez at the Wyss Institute for support with tissue culture. We thank Christophe Leterrier and Sebastian Strauss for helpful advice regarding optimal microtubule fixation. We thank Paul Guichard, Vincent Louvel, Kaibo Ma and Paul Tilberg for helpful discussions regarding fixation and hydrogels.

