## Supplementary Information for "DNA-PAINT imaging with hydrogel imprinting and clearing"


| | |
|---|---|
| **Supplementary Methods** | |
| **Supplementary Figure 1** | **Random selection of 256 DNA origami – Standard without hydrogel** |
| **Supplementary Figure 2** | **Random selection of 256 DNA origami – Hydrogel embedding** |
| **Supplementary Figure 3** | **Random selection of 256 DNA origami – Hydrogel embedding & clearing** |
| **Supplementary Figure 4** | **Hand-selected intact DNA origami – Hydrogel embedding & clearing** |
| **Supplementary Figure 5** | **Nearest-neighbor distance analysis for DNA origami data** |
| **Supplementary Table 1** | **Imaging parameters** |
| **Supplementary References** | |

### Supplementary methods

Materials

Unmodified, dye-labeled, and modified DNA oligonucleotides were purchased from Integrated DNA Technologies and Metabion. Unmodified oligos were purified via standard desalting and modified oligos via HPLC. DNA scaffold strands were purchased from Tilibit (p7249, identical to M13mp18). Sample chambers were ordered from Ibidi GmbH (8-well 80827). Tris 1M pH 8.0 (AM9856), EDTA 0.5M pH 8.0 (AM9261), Magnesium 1M (AM9530G) and Sodium Chloride 5M (AM9759) were ordered from Ambion. Streptavidin (S-888), ultrapure water (15568025), PBS (20012050), 4',6-Diamidino-2-Phenylindole, Dihydrochloride (D1306) (A39255), BSA (AM2616), DMEM (10569), Trypsin (25300-120), tetramethylethylendiamine (TEMED) (17919), ammonium persulfate (APS) (EN0531) and Dithiothreitol (DTT) were purchased from Thermo Fisher Scientific. BSA-Biotin (A8549), Tween-20 (P9416-50ML), Gelatin (G9391-100G), Glycerol (cat. 65516-500ml), (+-)-6-Hydroxy-2,5,7,8-tetra-methylchromane-2-carboxylic acid (Trolox) (238813-5G), methanol (32213-2.5L), 3,4-dihydroxybenzoic acid (PCA) (37580-25G-F), protocatechuate 3,4-dioxygenase pseudomonas (PCD) (P8279-25UN), Triton-X 100 (93443), 40% acrylamide (AA) (A4058), N,N'-Methylenebisacrylamide (BIS) Sigma-Aldrich (M1533-25ML), and poly-D-lysine solution (S2002) was purchased from Sigma-Aldrich. 10% fetal bovine serum was purchased from Genesee Scientific (25-514). 75 $cm^2$ cell flasks were purchased from Corning (353136). EM grade glutaraldehyde (GA) was purchased from Electron Microscopy Services (16220). 90 nm gold nanoparticles (G-90-20-10 OD10) were purchased from Cytodiagnostics. Primary anti a-tubulin (rat, MA1-80017) antibody was purchased from Thermo Fisher. Secondary donkey anti-rat and (711-005-152) antibodies were purchased from Jackson ImmunoResearch Laboratories. 0.5-mL Amino Ultra Centrifugal Filters with 50 kDa molecular weight cutoffs were purchased from Millipore (UFC5050). DBCO-sulfo-NHS ester cross-linker was purchased from Vector Laboratories (CCT-A124). Qubit Protein Assay (Q33211), NuPage 4-12% Bis-Tris protein gels (NP0323BOX), NuPage LDS Sample Buffer (NP0007) was purchased from Invitrogen. InstantBlue Coomassie Protein Stain was purchased from Abcam (ab119211).

Buffers

Four distinct buffers were used for sample preparation and imaging: Buffer A (10 mM Tris-HCl, pH 7.5, 100 mM NaCl); Buffer B (5 mM Tris-HCl, pH 8.0, 10 mM $MgCl_2$, 1 mM EDTA); Buffer C (1× PBS, 500 mM NaCl); and a 10× folding buffer (100 mM Tris, 10 mM EDTA, pH 8.0, 125 mM $MgCl_2$). Digestion buffer (50 mM Tris, 1 mM EDTA, 0.5% Triton X-100, 0.8 M guanidine HCl, pH 8)[1]. All buffers were checked for pH consistency stored at 4°C for several weeks. Prior to imaging, buffers were supplemented with an oxygen-scavenging and triplet-state quenching system containing 1× PCA, 1× PCD, and 1× Trolox.

PCA, PCD, Trolox system

100× Trolox: 100 mg Trolox was dissolved in 430 µL of 100% methanol, followed by the addition of 345 µL of 1 M NaOH and 3.2 mL of $H_2O$. 40× PCA: 154 mg PCA was dissolved in 10 mL of water and adjusted to pH 9.0 with NaOH. 100× PCD: 9.3 mg PCD was mixed with 13.3 mL of buffer containing 100 mM Tris-HCl (pH 8), 50 mM KCl, 1 mM EDTA, and 50% glycerol.

DNA origami design and assembly

DNA origami featuring 20-nm spaced adapter binding sites was previously designed using the Picasso Design[2] module. A complete list of the DNA strands used can be found in ref.[3]. The folding reaction was prepared with the following components: single-stranded DNA scaffold (0.01 µM), core staples (0.1 µM), biotin staples (0.01 µM), and extended staples for DNA-PAINT (each at 1 µM), all in a 1× folding buffer, with a total reaction volume of 50 µl per sample. Annealing was carried out by gradually cooling the mixture from 80 °C to 25 °C over 3h in a thermocycler. A 1:1 ratio of scaffold to biotin staples enabled sample preparation without prior DNA origami purification, as free biotinylated staples would otherwise saturate the streptavidin surface, preventing proper origami immobilization on the glass surface.

Oligonucleotide designs

The following DNA oligonucleotide sequences were used for all experiments: a 20-nt adapter binding site[4] (A20: AAGAAAGAAAAGAAGAAAAG), which allowed subsequent hybridization of desired docking strand to the origami via a complementary, stably binding adapter featuring the docking strand on the 5' and the acrydite moiety on the 3' end('docking-strand_cA20_acrydite': (CTCCTCCTCCTCCTC-tt-CTTTTCTTCTTTTCTTTCTT-acrydite). For microtubule immunolabeling, secondary antibodies were conjugated with the same A20 adapter binding site as previously used on DNA origami (see also "Conjugation of secondary antibodies"). A Cy3b-labeled imager strand was used for all experiments (Pm2: GAGGAGG-Cy3b).

Conjugation of secondary antibodies

DNA-antibody conjugation was carried out using 0.5-mL Amino Ultra Centrifugal Filters with a 50 kDa molecular weight cutoff. The reaction employed a DBCO-sulfo-NHS ester cross-linker, which was prepared at 20 mM in DMSO and stored in single-use aliquots at -80°C. This cross-linker facilitates covalent attachment of azide-functionalized DNA oligonucleotides to surface-exposed lysine residues on antibodies. The azide-functionalized DNA oligonucleotides were maintained in 1 mM deionized water. To ensure efficient conjugation, all antibodies were ordered in a carrier-free format, as preservatives like bovine serum albumin (BSA) and sodium azide interfere with the reaction. To begin, 500 µL of PBS was added to the Amicon filters and centrifuged at 10,000 rcf for 5 min to wet the membrane. Next, 25 µg of antibody was introduced and washed twice with PBS. For each wash, PBS was added up to 500 µL, followed by centrifugation at 10,000 rcf for 5 min. If the residual volume after the second wash exceeded 100 µL, an additional centrifugation step was performed under the same conditions. After washing, a 20-fold molar excess of DBCO-sulfo-NHS ester cross-linker and azide-functionalized DNA oligonucleotide was added to the antibody solution. The mixture was gently mixed and incubated overnight at 4°C in the dark to facilitate conjugation. The following day, the conjugated antibodies were washed three times with PBS using the same centrifugation steps described above. To elute the conjugated antibody, the filter was inverted into a fresh tube and centrifuged at 1,500 rcf for 2 min. The purified conjugate was then transferred to a clean tube and stored at -20°C in antibody storage buffer. Antibody concentrations were determined using the Qubit Protein Assay. To confirm successful DNA-antibody conjugation, unconjugated and conjugated antibodies were analyzed via NuPAGE 4–12% Bis-Tris protein gels. For each sample, 0.5 µg of total protein was mixed with NuPAGE LDS Sample Buffer and 50 mM DTT. Samples were denatured at 80°C for 10 min before gel electrophoresis, which was run at 75 V for 5 min, followed by 180 V for 60 min. Gels were stained with InstantBlue Coomassie Protein Stain for 15 min at room temperature, rinsed with water, and imaged using a Sapphire Biomolecular Imager (Azure Biosystems).

DNA origami sample preparation

Ibidi 8-well slides were prepared as follows. A 10 µl drop of biotin-labeled bovine albumin (1 mg/ml in buffer A) was placed at the center of the chamber, incubated for 2 min, and then aspirated. The chamber was subsequently washed with 200 µl of buffer A before aspirating again. Next, a 10 µl drop of streptavidin (0.5 mg/ml in buffer A) was added to the chamber center, incubated for 2 min, and aspirated. After washing sequentially with 200 µl of buffer A and 200 µl of buffer B, a 10 µl drop of DNA origami (1:100–200 dilution in buffer B from the folded stock) was introduced at the chamber center and incubated for 5 min. The chamber was then washed with 200 µl of buffer B, followed by the addition of docking strand adapters hybridizing to the DNA origami at 100 nM in buffer B. After a 5 min incubation, the chamber was washed with 200 µl of buffer B. Finally, buffer C and imager strand were added for DNA-PAINT imaging.

Cell culture

HEK293T were cultured in 75 $cm^2$ cell flasks at 37°C and 5%$CO_2$ in Dulbecco's Modified Eagle Medium (DMEM) with high glucose and Glutamax supplement, supplemented with 10% fetal bovine seralbumin. Cells were passaged once a week for no longer than 30 passages. For passaging, cells were washed with PBS, trypsinized, centrifuged, resuspended in fresh media, counted, and about 500 000 cells were transferred to a new flask with pre-warmed media. Aliquots of cells were stored in liquid nitrogen, in freezing medium (10% dimethylsulfoxide

[DMSO] in PBS) at a density of 1 million cells/ml. For imaging, cells were plated in Ibidi 8-well µ-Slides. For coating of glass surfaces, slides were incubated for 2 hours or overnight at 37°C with 200 µl of poly-D-lysine solution (diluted 1:10 in milliQ water) per well, rinsed twice with milliQ water and once with PBS. Cells were trypsinized, spun down, counted, and diluted to a concentration of 100 000 cells/ml. 200 µl or 2 ml of cells were placed in each chambered well and slides were incubated at 37°C and 5%$CO_2$ for 12-24 hours.

Cell fixation and immunolabeling

Our GA fixation protocol was adapted from those proposed by Jimenez et al.[5] and Strauss et al.[6]. Cells plated in slides or coverslips as described above were pre-fixed in 0.2% GA, 0.25% TritonX-100 in PBS for 90 seconds, fixed in 2%GA in PBS for 10 minutes at 37°C, rinsed three times in PBS, reduced in 0.1%w/v NaBH4 in PBS for 7 minutes, rinsed again three times in PBS and permeabilized in 0.25% TritonX-100 in PBS for 15 minutes. Both pre-fixative and fixative were pre-warmed to 37°C. For handling, slides were placed on aluminum block pre-warmed at 37°C, with a wet cloth between the slide and the block, and 200 µl of each solution were transferred to each well, one at the time. Coverslips were placed on plastic coverslip rack and moved to 100ml glass beakers containing the solutions. After permeabilization, samples were incubated with a blocking solution of 0.2% Gelatin and 0.1% TritonX-100 in PBS for 2 hours under gentle agitation. Primary rat anti-alpha tubulin monoclonal IgGs were diluted 1:100 in 0.2% Gelatin, and incubated in the wells for at least 4h at 4°C. After three 5-minute washes with 0.2% Gelatin 0.1% TritonX-100 in PBS, DNA-labelled secondary anti-rat polyclonal IgGs were incubated for 2 hours under gentle agitation in 0.2% Gelatin at a dilution of 1:100-1:200. protein low-bind Eppendorf tubes were always used to dilute antibodies. Volumes of 160 µl of antibodies and 200 µl of washing and blocking solutions were used for 8-well Ibidi slides. Finally, samples were washed 3× in PBS for 2min each and either processed for hydrogel imprinting or directly imaged in case of standard DNA-PAINT. Before imaging, samples were incubated with gold particles as fiducial markers (1:20 in PBS) for 5 min, washed again 2× in PBS before adding imaging buffer C.

Hydrogel imprinting and clearing

In general, we followed the hydrogel matrix imprinting and clearing concept proposed by Moffitt et al.[7] for anchoring acrydite-labeled oligonucleotides into non-expanding PA hydrogels consisting of acrylamide (AA), bis-acrylamide (BIS), TEMED and APS. However, we adapted the quantities of each component according to a more recent protocol for Ultrastructure Expansion Microscopy protocol (U-ExM) by Gambarotto et al.[8,9]. The following stock solutions were prepared and stored at -20°C: 500 µl aliquots of Monomer Solution (MS) consisting of 10% acrylamide (AA) and 0.1% BIS in 1x PBS; 50 µl aliquots of 10% TEMED in nuclease-free water; 50 µl aliquots of 10% APS in nuclease-free water. Prior to hydrogel imprinting, we added 100 nM acrydite-labeled docking strand adapter and incubated the samples for 10min for hybridization to DNA origami adapter binding sites/secondary antibodies (buffer B was used for DNA origami and buffer C for fixed cells). Samples rinsed 2x and washed in each buffer for 10 minutes. MS, TEMED and APS aliquots were thawed and kept on ice. 90 µl of MS were placed in a PCR tube. 5 µl of TEMED and 5 µl of APS were added in the tube and mixed by pipetting up and down for three times. 8-well slides were placed on ice, wells were filled with 200 µl of MS-TEMED-APS solution and incubated at 4°C for 15 minutes and then sealed and incubated at 37°C for one hour. Subsequently, samples were washed in 3x in PBS for 2 min. For hydrogel clearing, samples were incubated for a minimum of 12h of 1:10 Proteinase K diluted in digestion buffer followed by 3x washes in PBS for 2 min.

TIRF microscope

TIRF imaging was conducted at the MicRoN Imaging Core at Harvard Medical School using a Nikon Ti inverted microscope. The system was equipped with a Nikon Ti-TIRF-EM Motorized Illuminator, a Nikon LUN-F Laser Launch with a single fiber output (488 nm, 90 mW; 561 nm, 70 mW; 640 nm, 65 mW), and a Lumencore SpectraX LED Illumination unit. Imaging was performed using an oil-immersion objective lens (Apo TIRF 100×/1.49 DIC N2) within an objective-type TIRF system. For DNA-PAINT experiments, the 560 nm laser line was used for excitation, and fluorescence emission was filtered through a Chroma ZT 405/488/561/640 multi-band pass dichroic mirror and a Chroma ET 595/50m band-pass emission filter, before detection on a sCMOS camera (Andor Zyla 4.2) attached to a standard Nikon camera port.

Imaging conditions

All fluorescence microscopy data was recorded with the sCMOS camera (2048 × 2048 pixels, pixel size: 6.5 μm). Both microscope and camera were operated with the Nikon Elements software at 2×2 binning and cropped to the center 512 × 512 pixel field-of-view. The camera read out rate was set to 200 MHz and the dynamic range to 16 bit. A laser power of ~8 mW (measured behind the objective) was used for all data acquisitions and the TIRF coupling was adjusted prior to each experiment. All datasets were acquired for 5,000 frames at 200 ms exposure time. For additional imaging parameters and imager concentrations used see Supplementary Table 1.

Post-processing and image analysis

Raw fluorescence data was processed for super-resolution reconstruction using the 'Picasso' suite developed by the Jungmann lab[2]. Picasso Localize was used for spot-finding and saved as .hdf5 lists. The localization files were subsequently loaded into Picasso Render to perform drift correction (global correction via redundant-cross correlation[10] and subsequent correction based on fiducials). Picasso Render was used to export rendered images of super-resolved DNA origami structures and microtubule filaments in fixed cells and to calculate the NeNA localization precisions. Our clustering analysis to measure nearest neighbor distances between single docking strands on DNA origami nanostructures is described in Supplementary Fig. 5. Averaged DNA origami sum images were obtained by using the 'pick similar' tool in Picasso Render (default settings, pick diameter: 1px; range: 2.0) to select and export individual DNA origami structures from the data set and obtain averaged images using Picasso Average. The picking tool in Picasso Render was also used for measuring microtubule diameters (rectangular shape, width: 5px). A custom Python script was then used to calculate the width of the localization distribution (2x std.) with respect to the microtubule axis.

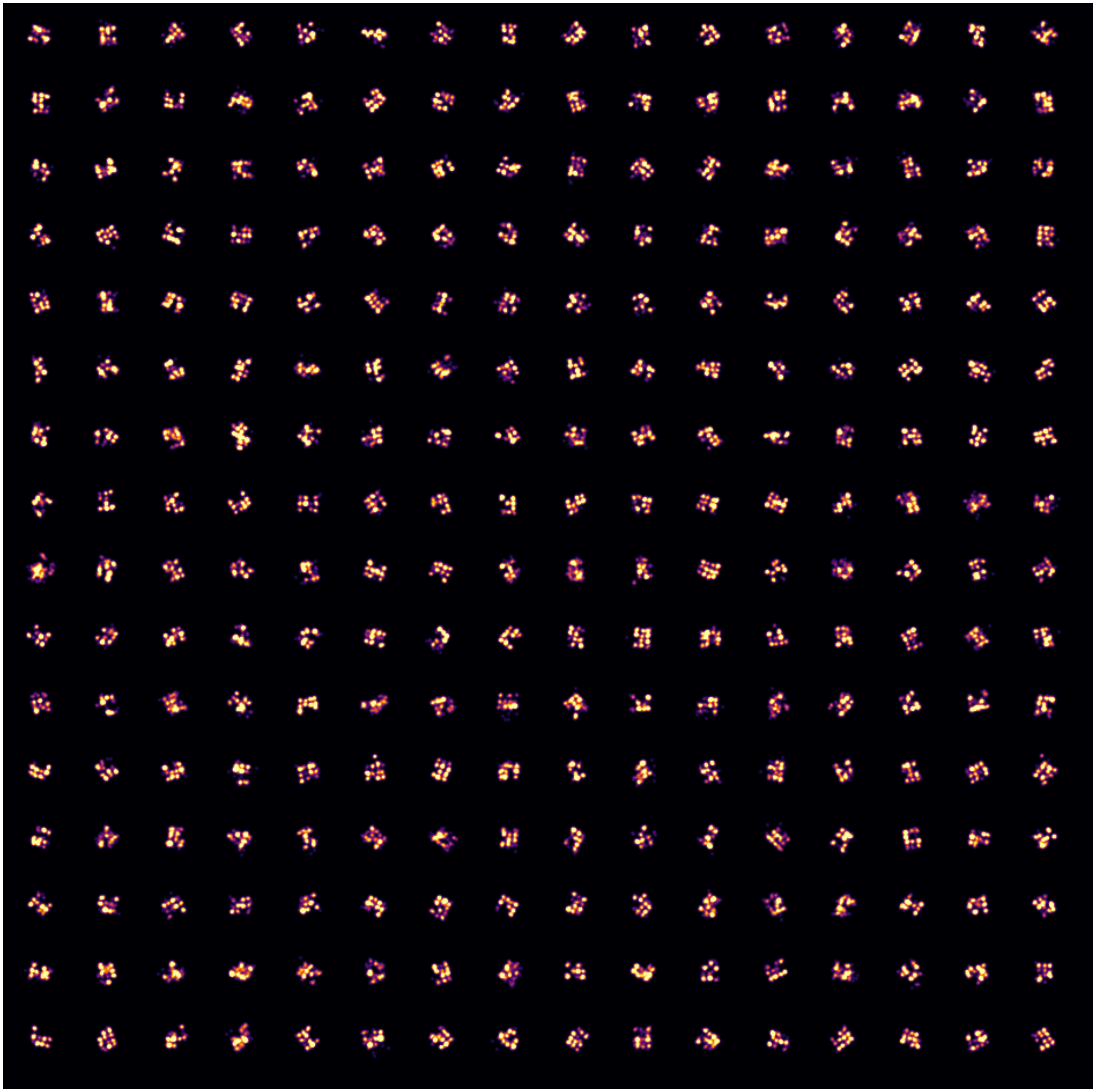

**Supplementary Figure 1**. Random selection of 256 DNA origami – Standard without hydrogel embedding. DNA origami structures are shown for the dataset used to obtain the averaged sum image of the 20 nm grid docking strand pattern in Fig. 1b.

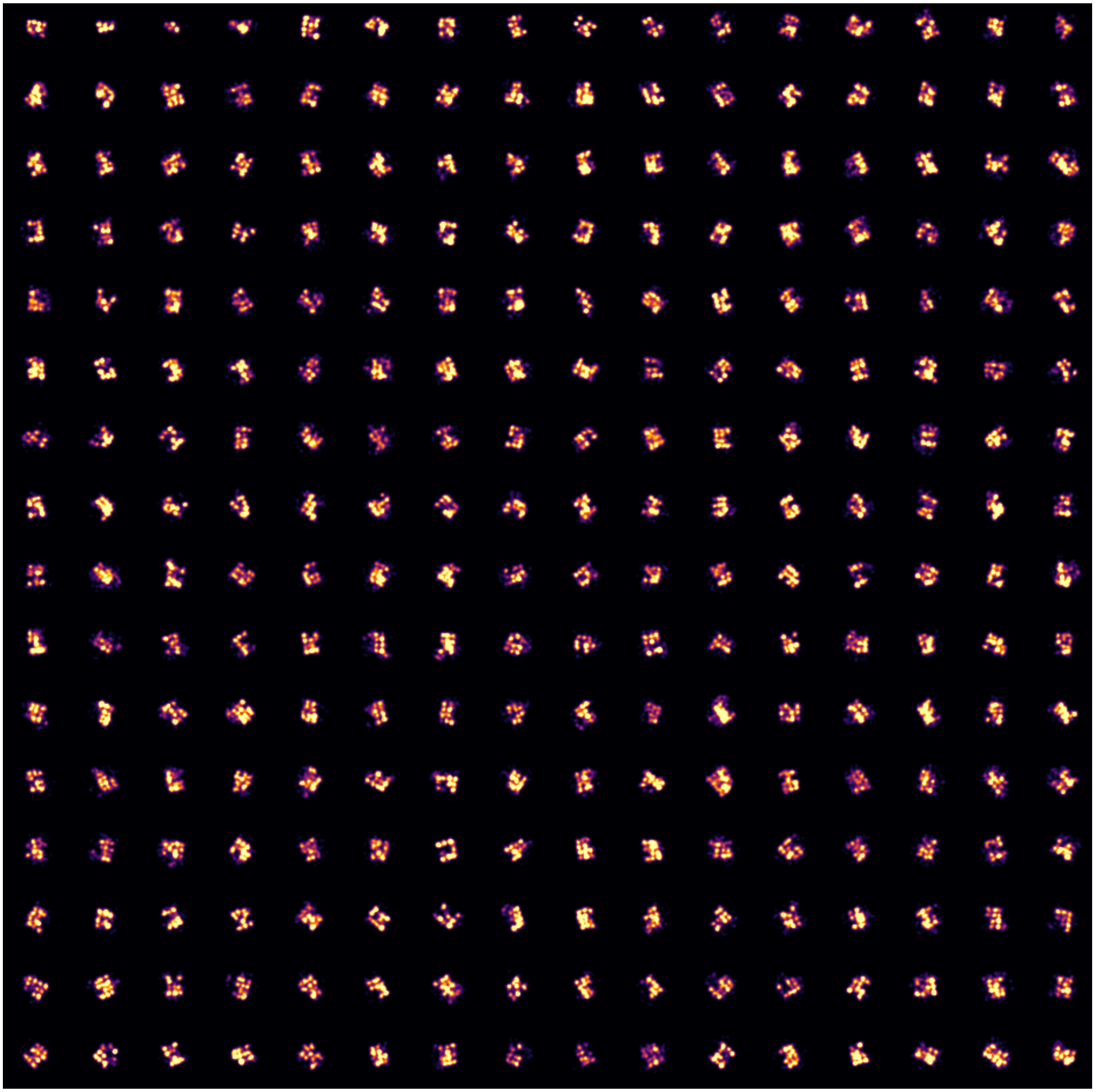

**Supplementary Figure 2**. Random selection of 256 DNA origami – Hydrogel embedding. DNA origami structures are shown for the dataset used to obtain the averaged sum image of the 20 nm grid docking strand pattern in Fig. 1c.

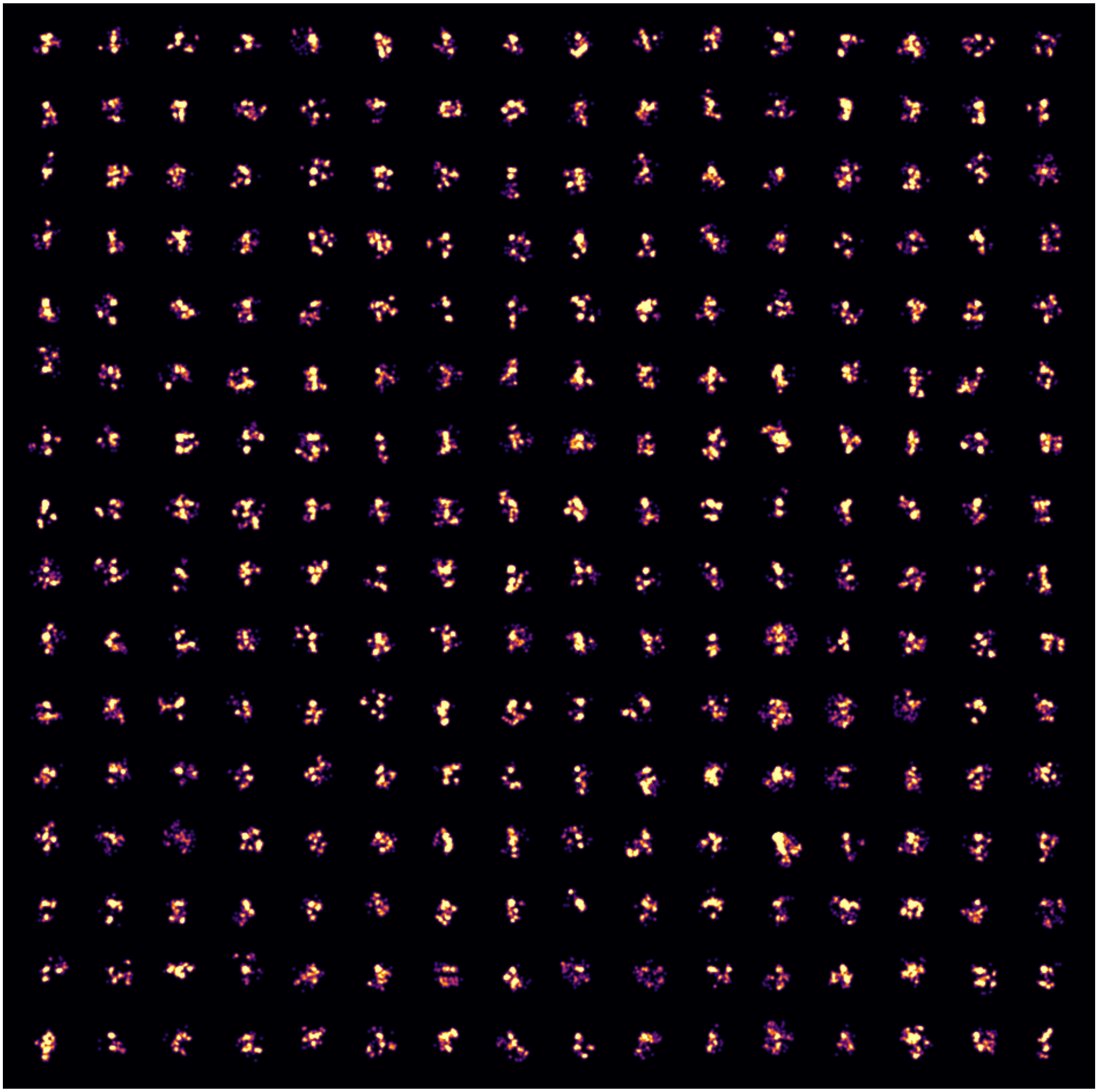

**Supplementary Figure 3**. Random selection of 256 DNA origami – Hydrogel embedding & clearing. After treatment with Proteinase K, individual DNA origami structures were still visible, but the 20 nm pattern was largely distorted. Since neither DNA nor polyacrylamide should be affected by Proteinase K activity, it is likely that prolonged incubation for 1 h at 37°C in low ion buffer led to structural deformation of DNA origami structures and their imprinted patterns at the hydrogel surface. Although few DNA origami patterns remained intact, random selection and averaging was not sufficient for reconstruction of the overall grid pattern.

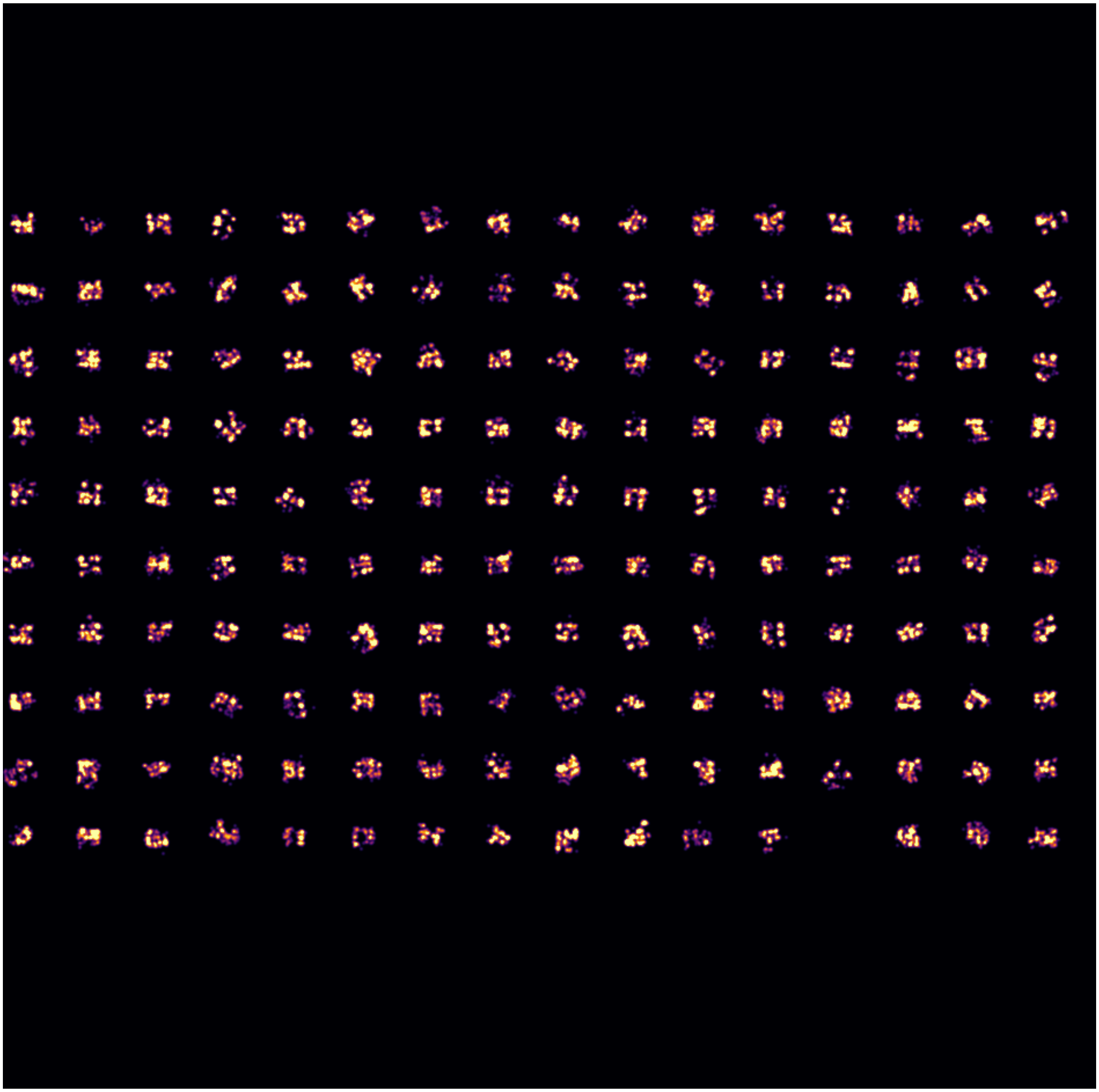

**Supplementary Figure 4**. Hand-selected intact DNA origami – Hydrogel embedding & clearing. To obtain the averaged DNA origami representation in Fig. 1d, we hand-selected intact DNA origami structures from the data set (n=159).

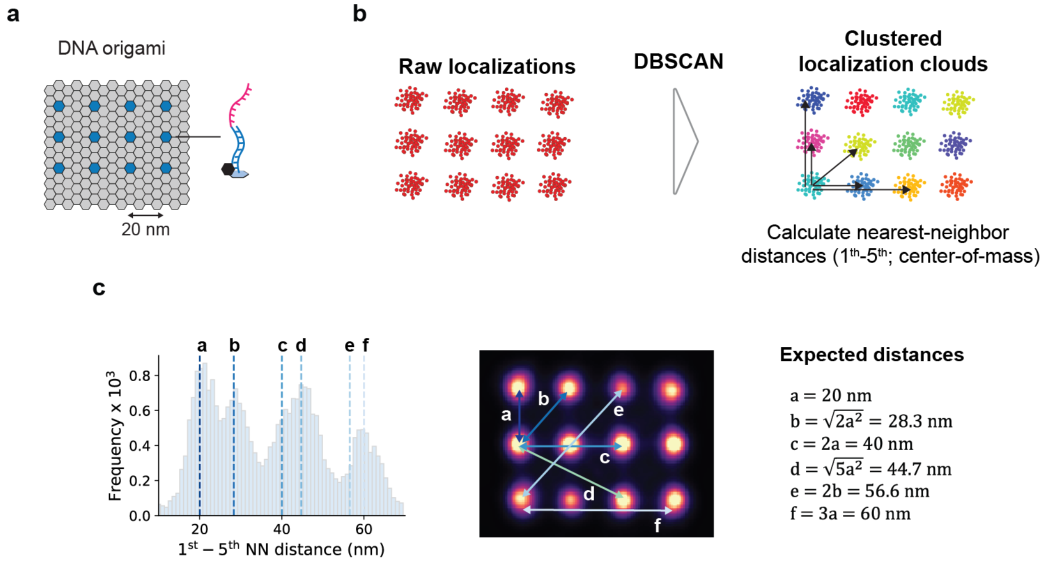


**Supplementary Figure 5**. Nearest neighbor distance analysis for DNA origami data. (a) Schematic of DNA origami nanostructures featuring a 3×4 pattern of binding sites for acrydite-labeled docking strand adapter oligonucleotides arranged at a 20 nm spacing. (b) Schematic of DBSCAN clustering analysis. A single DNA-PAINT docking strand is repeatedly visited by imager strands during data acquisition. Thus, localizations build up over time at each docking strand position, leading to accumulations of localizations referred to as localization clouds. The clustering algorithm DBSCAN[11] is a standard approach for localization cloud identification in DNA-PAINT[12] and implemented in the Picasso Render module[2]. Using *minPts*=6 (number of points required to form a dense region) and eps=$2x\sigma_{NeNA}$ (epsilon; distance between points to be considered as within neighborhood), we were able to identify single docking strand localization clouds on each DNA origami (each color represents an identified localization cloud). This allowed us to compute the center-of-mass distances for the 5 nearest neighbors for each localization cloud. (c) Nearest neighbor distances match the expected distance profile. Since docking strands are arranged at a 20 nm grid pattern, the expected distances can be calculated via geometry (a-f, blue arrows). The nearest neighbor distance profile on the left (here shown for a standard DNA-PAINT dataset without hydrogel-imprinting in Fig. 1e) highlights that peak positions match the expected distance profile, confirming our DBSCAN clustering parameter choice and overall structural integrity.

#### Supplementary Table 1 | Imaging parameters

| Figure | Sample | Condition | Imager concentration (nM) | Imaging Buffer | Exposure time (ms) | Frames |
|---|---|---|---|---|---|---|
| 1b<br>S1 | DNA origami | Standard (no hydrogel imprinting) | ~100 pM | B | 200 | 5,000 |
| 1c<br>S2 | DNA origami | With hydrogel imprinting | ~500 pM | B | 200 | 5,000 |
| 1d<br>S3-4 | DNA origami | With hydrogel imprinting & clearing | ~500 pM | B | 200 | 5,000 |
| 2b,e-f | HEK cells | Standard (no hydrogel imprinting) | ~50 pM | C | 200 | 5,000 |
| 2c,e-f | HEK cells | With hydrogel imprinting | ~500 pM | C | 200 | 5,000 |
| 2d,e-f | HEK cells | With hydrogel imprinting | ~50 pM | C | 200 | 5,000 |
